# Single-molecule tracking in the psychrophile *Psychrobacter cryohalolentis* reveals heat-shock-induced changes in ClpX dynamics

**DOI:** 10.64898/2026.11.16.767012

**Authors:** Eveliny Tomás Nery, Yuan Sui, Jarno Mäkelä

## Abstract

Psychrophilic organisms display a series of physiological adaptations which enable them to thrive in cold environments, yet the dynamics of these processes in living cells remain largely unexplored. Here, we establish super-resolution microscopy and single-molecule tracking (SMT) in the cold-adapted bacterium *Psychrobacter cryohalolentis*, enabling the investigation of protein dynamics *in vivo* at high spatiotemporal resolution. For controlled expression of HaloTag fusions in this organism, we developed an inducible plasmid expression system based on a galactose-inducible promoter. As a proof of concept, we followed the AAA+ unfoldase ClpX during steady-state growth and heat shock. At 22 °C, ClpX diffused as a single population with a media apparent diffusion coefficient of 0.25 µm²/s, suggesting that most molecules reside in large complexes. A shift to 32 °C, above the growth maximum of this strain, broadened the mobility distribution, increasing both slow and fast populations, and transiently redistributed ClpX towards the cell periphery. During recovery the fast population returned to baseline within 5 min, whereas the slow population remained elevated for at least 30 min, suggesting that the two populations reflect distinct processes. SMT in a psychrophilic bacterium thus opens new possibilities to study how cold-adapted microorganisms respond to environmental change at the molecular level.

## Introduction

Although model organisms such as *Escherichia coli* have served as powerful systems for studying molecular dynamics, they represent only a small fraction of the environmental conditions under which life can thrive, which limits our understanding of the full diversity of biological adaptations. This overlooked diversity includes psychrophiles (here used broadly for cold-adapted organisms, including psychrotrophs with growth optima above 15 °C), microorganisms capable of growth and metabolic activity at temperatures that are inhibitory or lethal for most organisms. To grow in these habitats, psychrophiles have evolved molecular and physiological adaptations, such as cold-active enzymes, membranes of higher fluidity, and specialized stress-response systems, that allow them to maintain cellular function at low temperature, and often also under high salinity and limited nutrient availability [1, 2, 3, 4]. These adaptations also underpin their biotechnological value, as cold-active enzymes are widely exploited in applications ranging from detergent formulation and food processing to pharmaceutical production and waste degradation [5, 6, 7, 8, 9, 10, 11]. One example of such an organism is *Psychrobacter cryohalolentis* K5, a Gram-negative bacterium isolated from a Siberian permafrost cryopeg that grows between −10 and 30 °C [12]. Its wide thermal range therefore spans both psychrophilic and mesophilic conditions within a single strain, making it an attractive model for investigating the molecular basis of cold adaptation and temperature stress.

A central process for temperature adaptation is proteostasis, which maintains the cellular proteome through the folding, trafficking and degradation of misfolded, damaged, or unnecessary proteins [13]. Proteostasis is expected to be particularly demanding in psychrophiles, whose proteins are generally more flexible and less thermally stable than their mesophilic counterparts [14, 15], so that temperatures tolerated easily by a mesophile may already impose proteotoxic stress. In *E. coli*, ClpX is an ATP-dependent AAA+ translocase and unfoldase protein that acts as the substrate-specificity component of the ClpXP protease and can also function as a chaperone on its own [16]. It recognises proteins carrying the 11-residue SsrA tag, added to their C-terminus when ribosomes stall [17], and uses cycles of ATP binding and hydrolysis to unfold these substrates and pass them into the ClpP peptidase for degradation [17, 18]. ClpX expression is induced during heat shock and has been implicated in stress tolerance in several bacteria [19, 20]. However, it remains unclear how ClpXP behaves during thermal stress in an organism whose proteome is adapted to function at low temperature.

In mesophilic organisms, single-molecule tracking (SMT) fluorescence microscopy has emerged as a powerful tool for investigating protein dynamics directly within living cells, enabling the observation of individual molecules with high spatial and temporal resolution [21, 22]. By following the motion of fluorescently labelled proteins over time, SMT quantifies their diffusion behaviour and allows functional interactions to be inferred, such as binding to DNA, ribosomes, or other cellular structures, that would be obscured in ensemble-averaged measurements [21, 23, 24]. The approach has been applied in model organisms such as *E. coli* to characterize proteins including ribosomal subunits, transcription factors and chaperones, providing insight into their *in vivo* behaviour under different environmental conditions [25, 26, 27, 28, 29]. More recently, SMT has been extended to the halophilic archaeon *Haloferax volcanii*, where it has resolved the dynamics of RNA polymerase, the division protein FtsZ1, and the response of the CRISPR-associated factor Cas1 to UV-induced DNA damage [30]. To our knowledge, however, SMT has not been applied in any psychrophile.

Here, we establish single-molecule tracking in *P. cryohalolentis* K5, enabling the direct investigation of protein dynamics *in vivo* at high spatiotemporal resolution. We first developed an inducible expression system for this organism, based on the galactose-inducible promoter of another psychrophile, *Pseudoalteromonas translucida* TAC 125 [31], allowing controlled expression of tagged proteins. We then applied this platform to characterize the dynamics of the AAA+ translocase and unfoldase ClpX during steady-state growth and heat shock, providing insight into how the protein quality control machinery of a psychrophilic bacterium responds to thermal stress at single-molecule resolution.

## Materials and Methods

### Strains and growth conditions

Wild-type strains *of P. cryohalolentis* K5 (DSM 17306) and *Pseudoaltermonas translucida* TAC125 (CIP 108707; formerly *P. haloplanktis*) were obtained from the German Collection of Microorganisms and Cell Cultures (DSMZ, Braunschweig, Germany) and the Collection of the Institut Pasteur (CIP, Paris, France), respectively. Both strains were grown in Marine Broth (MB, Difco™ 2216) at 22 °C in a rotary shaker (220 rpm). For solid media, agar was added at a final concentration of 1.5% (w/v). *Escherichia coli* strain DH5α was cultured in Lysogeny Broth (LB; Sigma-Aldrich) at 37 °C with shaking. Where required, kanamycin (kan) was added to culture medium at the final concentration of 25 and 50 μg/ml for *P. cryohalolentis* and *E. coli*, respectively.

### Construction of expression plasmid (pPS-BR_gal)

A vector for inducible gene expression was constructed using the galactose operon promoter (galP) from *P. translucida* [31] and the shuttle vector pPS-BR [32] as backbone. The chromosomes were extracted using the Zymo *Ǫuick-*DNA Miniprep kit (Zymo Research). Ǫ5® High-Fidelity DNA Polymerase (New England Biolabs) was used in all PCRs. GalP had been previously amplified from the genome of *P. translucida* using primers gal.F/R (Table S1) and cloned into vector pSEVA221 [33] between restriction sites XbaI and PacI producing the construct pSEVA221_gal. For this study, this insert was amplified from pSEVA221_gal along with the vector’s original λ transcription terminator (T_0_) using primers seva.gal.F/R (Table S1) and the backbone of pPS-BR was amplified using primers ppsbr.F/R (Table S1). After that, both amplicons were treated with DpnI restriction endonuclease (New England Biolabs) and assembled using the NEBuilder HiFi DNA Assembly Master Mix (New England Biolabs) generating pPS-BR_gal. This construct was then transformed into chemically competent *E. coli* DH5α cells. Presence of the insert containing the galactose promoter was checked by colony PCR using the pair of primers BR-insert.F/R. The plasmid was purified from a positive colony using ǪIAprep Spin Miniprep Kit (ǪIAGEN) and the assembly was further confirmed by whole-plasmid Sanger sequencing (Eurofins Genomics, Ebersberg, Germany).

### Construction of HaloTag fusion plasmid (pPS-BR_gal-Halo)

Using PPS-BR_gal as backbone, we assembled a construct bearing the sequence coding for HaloTag7 with a linker (VDGSAGSAAGSGEF) for producing protein-HaloTag C-terminal fusions. This sequence was amplified from pSU005 [34] using primers Halo-link.F/R (Table S1) which were designed with SalI and PstI restriction sites. The amplicon was PCR cleaned using the ǪIAquick PCR Purification Kit (ǪIAGEN) and digested with SalI and PstI restriction enzymes (New England Biolabs). This fragment was then ligated into pPS-BR_gal using T4 ligase (New England Biolabs) and the resulting construct was transformed into *E. coli* DH5α cells. The resulting colonies were checked by colony PCR for the insert using the pair of primers BR-insert.F/R. The final construct was named pPS-BR_gal-Halo.

### Construction of ClpX-HaloTag plasmid (pPS-BR_gal-Halo_ClpX)

For obtaining the ClpX-Halotag (ClpX-HT) fusion plasmid, the *ClpX* gene was amplified from the chromosome of *P. cryohalolentis* without the stop codon and with AvrII and SalI restriction sites using the pair of primers ClpX.F/R. This amplicon was then cleaned using the ǪIAquick PCR Purification Kit (ǪIAGEN) and, along with pPS-BR_gal-Halo backbone, digested with AvrII and SalI restriction enzymes (New England Biolabs). The digested DNA fragments were ligated into pPS-BR_gal_Halo using T4 ligase (New England Biolabs) generating the plasmid pPS-BR_gal-Halo_ClpX. The resulting construct was transformed into *E. coli* DH5α cells and the presence of the ClpX genes in the resulting colonies was checked by colony PCR using primers BR-insert.R and BR-Halo-insert.F. This plasmid and all the other ones created in the present study were purified using the ǪIAprep Spin Miniprep Kit (ǪIAGEN) and confirmed by Sanger sequencing (Eurofins Genomics, Ebersberg, Germany). Previous work describes functional C-terminal ClpX fluorescent protein fusions [35, 36, 37].

### Transformation into *P. cryohalolentis*

pPS-BR_gal-Halo_*ClpX* and pPS-BR_gal (empty plasmid) were delivered to *P. cryohalolentis* by electroporation. In brief, the strain was inoculated in Marine Broth (MB; Marine Broth 2216, Millipore®) and incubated in a rotary shaker at 22 °C until reaching the late-exponential growth phase. Next, 5 mL of bacterial culture was cooled in ice for 15 min, centrifuged, and washed with 1 ml of cold sterile milli-Ǫ water three times to remove traces of the medium. The cells were then finally resuspended in 100 µL of cold milli-Ǫ water containing approximately 1 µg of plasmid and placed into an electroporation cuvette (1 mm gap; Fisherbrand). The electroporation was carried using a Gene Pulser XcellTM electroporator (Bio-Rad Laboratories) configured to deliver a pulse with field strength of 20 kV/cm (2000 V, 200 Ω, 25 µF). Immediately after the pulse, the cells were resuspended in 1 mL of MB supplemented with 2% NaCl (pre-warmed to 22 °C) and allowed to recover for 18-20 h in a rotary shaker at 22 °C. After that, the cells were spread in Marine Agar (MA) supplemented with 25ug/ml of kan and incubated under the same conditions until colonies could be observed (approximately 3 days).

### HaloTag expression and labelling

Strains transformed with pPS-BR_gal-Halo_*ClpX* or pPS-BR_gal were inoculated in MB supplemented with 25 μg/ml kan and incubated overnight at 22 °C in a rotary shaker (220 rpm). On the next day the cultures were diluted and incubated overnight in the same conditions until cells reached mid-exponential growth phase (OD600 = 0.15-0.25). When the desired OD600 was reached, pPS-BR_gal-Halo_*ClpX* bacterial cultures were treated with 0 (untreated), 0.25, 0.5 or 1% (w/v) galactose and incubated for 1 h along with untreated pPS-BR_gal strains.

For fluorescence labelling, the bacterial cultures were pelleted and resuspended in 100 μl MB to which Janelia Fluor 552 (JF552) dye was added to a final concentration of 0.5 μM. The tubes were then incubated at 22 °C for 30 min with shaking to label HaloTag. After that period, to remove the excess of dye, cells were washed five times, resuspended in pre-warmed MB and allowed to recover for 1 h at 22 °C with shaking. This recovery step allows the unbound dye inside the cells to diffuse out, diminishing background fluorescence. After recovery, each sample was pelleted, resuspended in PBS, added to 96 well plates (Thermo Scientific™) for absorbance (at 600 nm) and fluorescence (excitation at 539 nm and emission at 580 nm) measurements (Agilent BioTek Synergy H1 Multimode Reader). Normalized fluorescence intensity (RFU/OD600) was compared across all conditions (empty-vector control and each galactose concentration) using a one-way ANOVA followed by Tukey’s HSD post-hoc test for pairwise comparisons.

### Single-Molecule Imaging of ClpX

Bacterial cultures of *P. cryohalolentis* with pPS-BR_gal-Halo_*ClpX* were induced with 0.5% galactose and labelled as above. After that, the cells were spotted over a pre-warmed 1% gellan gum (Thermo Scientific Alfa Aesar) pad prepared with MB and allowed to dry at 22 °C. The pad was then placed on a VaHEAT chip (Stable Scopes) which had been previously plasma-etched (Tergeo plasma cleaner; Pie Scientific) to remove background fluorescent particles. Sample temperature was controlled and monitored during imaging using the VaHEAT heating stage.

Live-cell single molecule localization microscopy and single molecule tracking was performed on a custom TIRF microscope built around a Rapid Automated Modular Microscope (RAMM) System (ASI Imaging) equipped with a motorized stage, a z-motor objective mount, and a motorized filter wheel (US-2000, MIM4, TG8; ASI Imaging). Fluorescence excitation was provided by a 561 nm laser (100 mW, 50% transmission; iChrome MLE, Toptica). The laser was collimated and focused through a 100x oil-immersion objective (CFI Plan Apochromat Lambda D, NA 1.45; Nikon) and directed onto the sample at an angle to achieve highly inclined and laminated optical sheet (HILO) illumination [38]. Fluorescence emission was separated from excitation light using a dichroic mirror and notch filter (ZT405/488/561rpc and ZET405/488/561m; Chroma) and detected with an EMCCD camera (iXon Life 897, 512 x 512 pixels; Andor). LED Illumination and condenser (ASI imaging) were used for acquiring phase contrast images for cell segmentation. Movies were acquired at a frame time of 7.475 ms, with 20,000 frames collected per acquisition.

For heat shock experiments, cells were first imaged at a baseline temperature of 22 °C and after that, the temperature was shifted to 32 °C. Cells were imaged at 5 and 15 min after 32 °C was reached and the temperature was then shifted back to 22 °C. Cells were imaged during recovery at 5, 15, and 30 min at 22 °C. The experiment was performed in three biological replicates.

### Single molecule tracking analysis

All analyses described in this study were performed in Python using NumPy 1.26.4, pandas 2.2.3, SciPy 1.13.0 and Matplotlib 3.10.3. Single molecule tracking was performed as in [25]. Fluorescent spot detection and tracking were performed using Trackpy [39]. For each frame of the time-lapse movie, fluorescent spots were detected using band-pass filtering, peak finding and intensity thresholding. These initial localizations were then used as starting points for high-resolution position refinement. Within cell areas, localizations in consecutive frames that were within 0.4 µm of each other were linked into tracks, allowing one missed frame within a track. Steps larger than 0.4 µm were considered erroneous as above this value, the step-size distribution forms a flat tail (Fig. S1a, b). If multiple localizations were detected within the detection radius, these localizations were discarded from the analysis. Tracks containing fewer than 9 displacements were omitted due to lower certainty of the calculated apparent diffusion coefficients (D*_app_*). Cell areas were generated from phase contrast images using Omnipose [40] for cell segmentation. The model “bact_phase_omni” was used for segmentation of our data and cells with bright immobile spots were excluded from analysis at this stage.

The mobility of molecules was determined by calculating the apparent diffusion coefficient from the stepwise mean-squared displacement using:

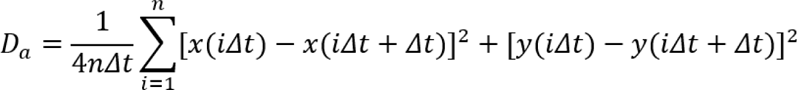

where *n* is the number of displacements in a track, *Δt* is the time between frames, and *x(iΔt)* and *y(iΔt)* are the coordinates of the molecule at frame *i* [24].

For each condition, tracks from all biological replicates were pooled, with one D*_app_* value per track. log₁₀ D*_app_* were plotted as probability density histograms (bin width 0.1 log₁₀ units) and estimated with a Gaussian kernel density estimate, using the same bandwidth for all conditions (0.11 log₁₀ units). The 22 °C kernel density estimate was subtracted from each heat-shock and recovery condition, on a common grid of log₁₀ D*_app_* values. Because both distributions are normalized, positive values indicate D*_app_* ranges over-represented in the condition and negative values ranges over-represented at 22 °C. The fraction of tracks that shifted between D*_app_* ranges relative to 22 °C was calculated as half of the integrated absolute difference.

### Single-molecule localization analysis

For each cell, the major (length; PC0) and minor (width; PC1) axes were found by principal component analysis of the pixel coordinates of the cell mask. Localizations were projected onto these axes and normalized by the cell half-length. Because the two cell poles cannot be distinguished, the major axis coordinate was folded. Cells with a projected area ≥ 2.5 µm² were excluded to avoid merged or dividing cells. Localizations from all cells measured in each condition were pooled into a 2D histogram (bin 0.08 units) as function of percentage of all localizations falling in each bin (colour scale 0–2.5%). The cell outline was calculated from the masks in the same normalized coordinates.

For the 1D plots, localization density along each axis is shown as a 1D histogram with a Gaussian kernel density estimate (Scott’s bandwidth × 0.63). To show the uniform localization profiles, the cell was modelled as a cylinder of radius R with two hemispherical caps (spherocylinder), set to the mean values measured from the cell masks at 22 °C. The expected profile across the major axis (PC0) is proportional to the cross-sectional area at each position. Across the minor axis (PC1), the profile is proportional to the area of the cell section at each position, [2L√(R² − y²) + π(R² − y²)]/V, where L is the cylinder length and V the cell volume. The reference profiles were overlaid on the observed distributions, so deviations from them show enrichment or depletion relative to an even distribution.

## Results

### Establishing galactose induction system in *P. cryohalolentis*

To express fusion proteins for single-molecule imaging in *P. cryohalolentis*, we cloned the galactose-inducible promoter galP from *Pseudoalteromonas translucida* [31] into the shuttle vector pPS-BR [32] and added a linker followed by HaloTag7 (HT), creating a general fusion platform (Fig. 1a). Although *P. cryohalolentis* and *P. translucida* belong to different orders within Gammaproteobacteria, their conserved transcriptional and translational machinery makes heterologous promoters likely to remain functional. As a proof of principle, we fused HT to the AAA+ unfoldase ClpX (ClpX-HT) and transformed the resulting plasmid into *P. cryohalolentis*. Because ClpX is a hexamer [16, 17, 18] and the chromosomal *clpX* gene was not deleted, ClpX-HT is expected to assemble into mixed hexamers containing both native and tagged subunits (Fig. 1b).

**Figure 1.**
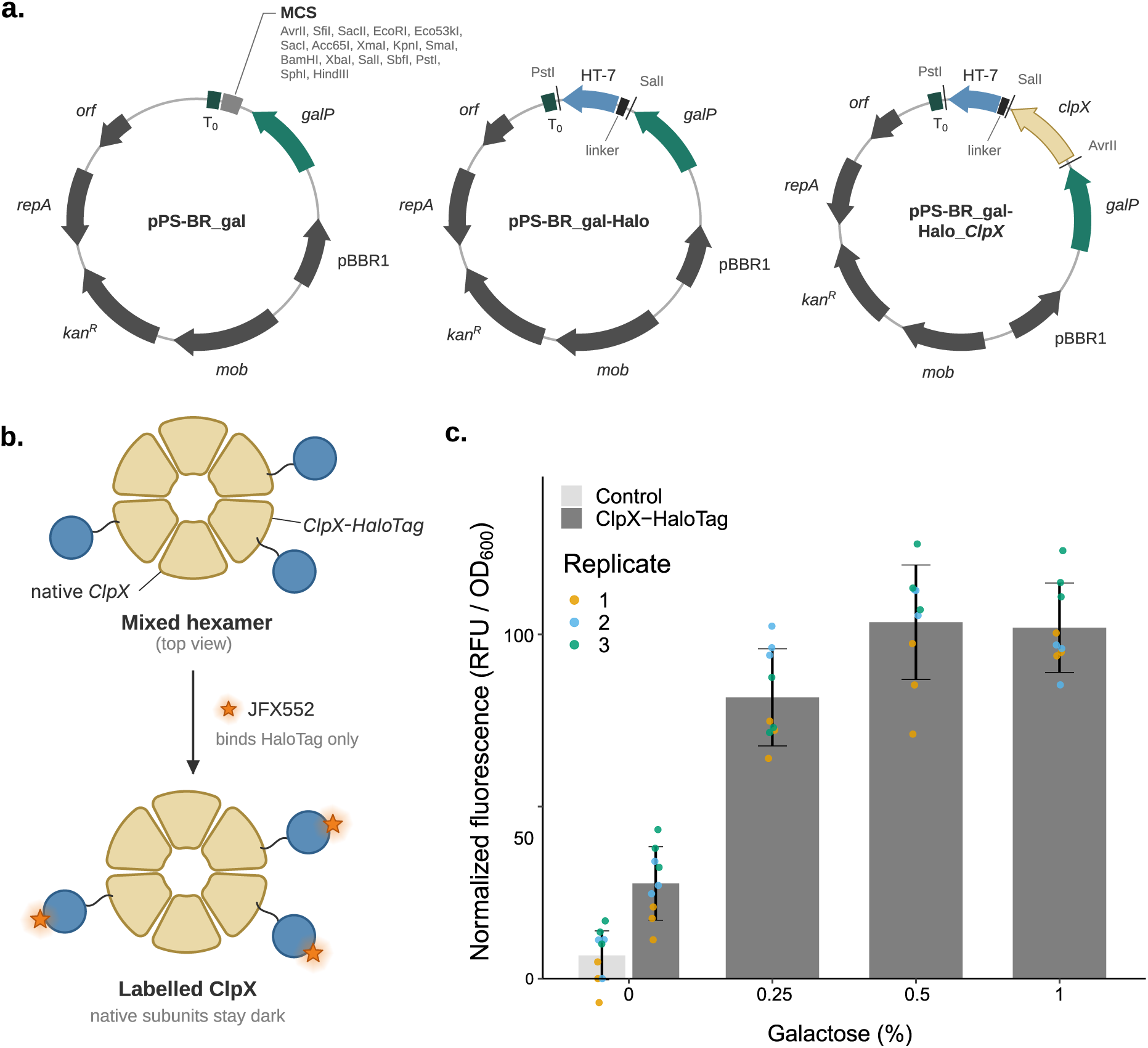
A galactose-inducible HaloTag expression system for P. cryohalolentis. (**a**) Genetic organization of the shuttle vectors pPS-BR_gal, pPS-BR_gal-Halo and pPS-BR_gal-Halo_ClpX. mob, relaxase gene for conjugal transfer; repA, replication initiation protein of the pP32BP2 from P. cryohalolentis; kan^R^, resistance gene aphA; pBBR1, replication initiation protein for replication in E. coli; orf, conserved hypothetical protein of plasmid pP32BP2; T_0_, strong Rho-independent terminator from bacteriophage lambda; MCS, multiple cloning site; HT-7, HaloTag7; (**b**) Schematic of the ClpX hexamer assembled from native and HaloTag-fused subunits. HT-7-fused subunits are labelled with JF552, rendering the complex visible by ffuorescence microscopy; (**c**) Normalized ffuorescence of P. cryohalolentis cultures carrying pPS-BR_gal (empty-vector control) and pPS-BR_gal-Halo_ClpX (ClpX-HT), induced with 0, 0.25, 0.5 or 1% galactose and labelled with JF552 dye. Bars show mean ± SD of three biological replicates. One-way ANOVA, F = 36.7, p = 5.99×10⁻⁶; p-values from Tukey’s post-hoc test.

To test whether the galactose-inducible system was suitable for ClpX-HT expression in *P. cryohalolentis*, we induced cultures with a range of galactose concentrations, labelled HT with the cell-permeable dye Janelia Fluor 552 (JF552) and measured total culture fluorescence (Figure 1c). Cultures carrying pPS-BR_gal without cargo served as a control for autofluorescence. Fluorescence increased from 27.7 ± 10.7 RFU/OD600 in non-induced cells to 81.7 ± 14.1 RFU/OD600 with 0.25% galactose and plateaued at 104.0 ± 16.6 and 102 ± 13 RFU/OD600 with 0.5% and 1% galactose, respectively (mean ± SD, n = 3 biological replicates; Figure 1c). All induced cultures differed significantly from both the empty-vector control and non-induced cells (p ≤ 0.003), while 0.5% and 1% galactose were indistinguishable (p > 0.99; one-way ANOVA with Tukey’s post-hoc test). Expression therefore saturates at 0.5% galactose, which we used in all subsequent experiments.

Non-induced ClpX-HT cultures showed about four-fold higher fluorescence than the empty-vector control (27.7 ± 10.7 vs. 6.8 ± 7.1 RFU/OD600, mean ± SD, n = 3), although this difference was not significant after correction for multiple comparisons (Tukey HSD, p = 0.33). With three replicates the comparison is underpowered, and the trend is consistent with low-level basal expression, arising either from trace galactose in the MB medium or from promoter leakage. Taken together, the galactose-inducible system is functional in *P. cryohalolentis*, driving dose-dependent expression of the HT fusion that saturates at 0.5% galactose in cells grown aerobically at 22 °C in MB.

### ClpX forms a single, uniformly distributed population at 22 °C

To establish a baseline for ClpX dynamics in *P. cryohalolentis*, we tracked ClpX-HT in cells grown at 22 °C. Cells were labelled sparsely with JF552 so that individual molecules could be localized in consecutive frames and linked into trajectories (Fig. 2a), and an apparent diffusion coefficient (D*_app_*) was calculated from the mean squared displacement of each trajectory. The distribution of D*_app_* was unimodal and close to symmetric (Fig. 2b), with a median of 0.25 µm²/s and an interquartile range of 0.16 to 0.37 µm²/s). Only 7.5% of trajectories fell below 0.10 µm²/s and 8.2% above 0.6 µm²/s, indicating that most ClpX molecules belong to a single population, with no separated immobile or freely diffusing sub-populations. Given the known relationship between protein size and cytoplasmic diffusion, below that expected for a large cytoplasmic complex such as the ClpX hexamer, alone or bound to ClpP or to substrates [41, 42].

**Figure 2.**
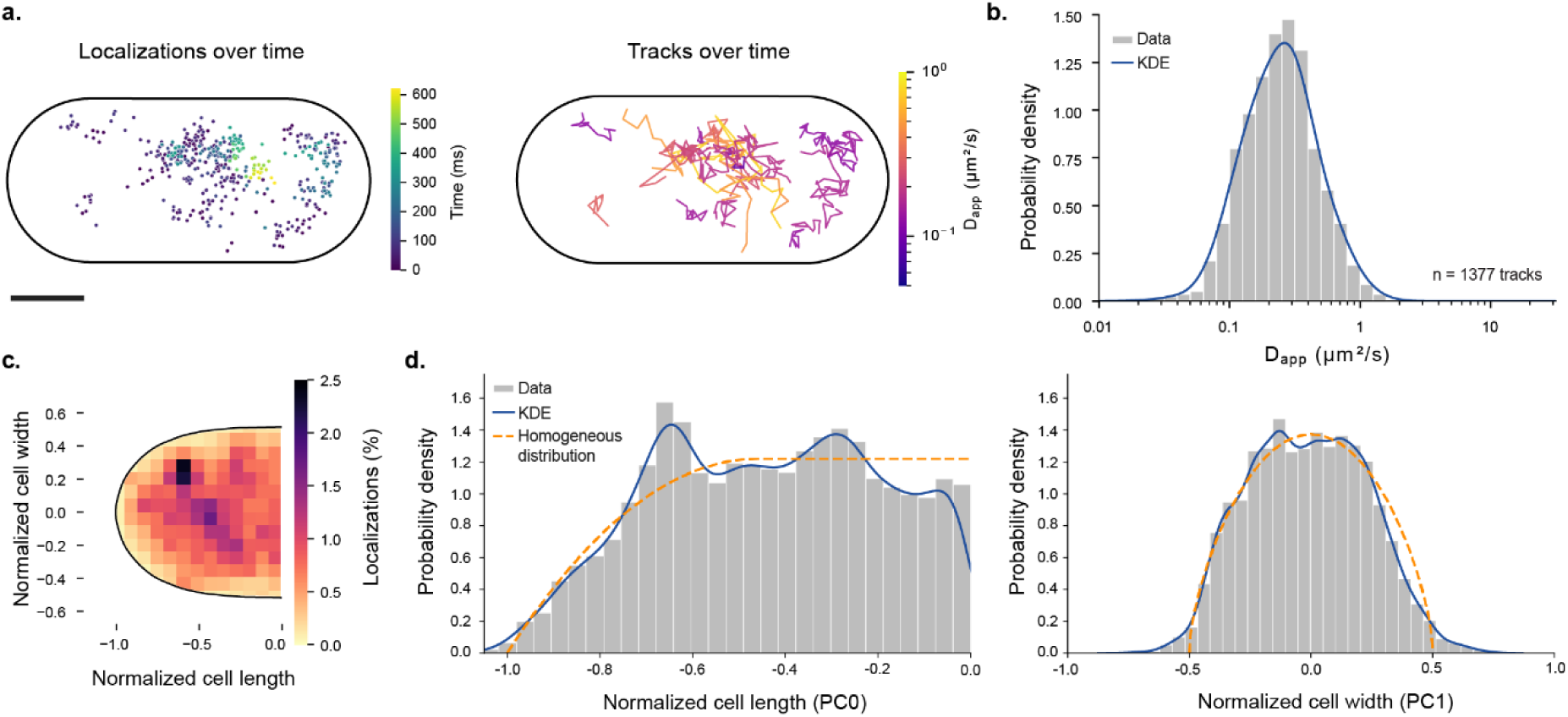
Single-molecule tracking of ClpX-HT in P. cryohalolentis at 22 °C. (**a**) Localizations of JF552-labelled ClpX-HT recorded over consecutive frames for a representative cell (left). Trajectories coloured by apparent diffusion coefficient (D_app_, log-scale) (right). Black line, cell outline. Scale bar, 0.5 µm. (**b**) Distribution of D_app_ (log-scale) for trajectories with at least S displacements. Blue line, Gaussian kernel density estimate (KDE; bandwidth 0.11 log₁₀ units). n = 1,377 trajectories from 251 cells. (**c**) Localization map in normalized, folded cell coordinates (pole at −1, mid-cell at 0), coloured by the percentage of localizations per bin. Black line, cell outline. n = 22,88S localizations from 251 cells. (**d**) Distribution of localizations along the long axis (PC0, left) and short axis (PC1, right). Blue lines, KDE (Scott’s bandwidth × 0.C3). Orange dashed lines, the profile expected for molecules distributed evenly through the volume of a rod-shaped cell (a cylinder with hemispherical caps). All data from three biological replicates.

At this condition, ClpX-HT was found distributed throughout the cytoplasm (Fig. 2c, d). Along the long axis, the localization density broadly followed the homogeneous reference, with small local maxima at about −0.65 and −0.3 and slight depletion close to the pole (Fig. 2d, left) while no polar enrichment was observed. Note that the apparent drop at mid-cell (position 0) is expected, since the profile is computed by folding each cell about its midpoint. Across the short axis, the profile closely matched the homogeneous reference (Fig. 2d, right), indicating no preference for the membrane or the cell centre. The single high-density bin in the localization map (−0.6, +0.25) most likely reflects repeated random sampling of a few molecules. Taken together, at 22 °C ClpX-HT in *P. cryohalolentis* diffuses as a single population and is evenly distributed in the cytoplasm, providing the baseline for the heat-shock and recovery experiments described in the next section.

### Heat shock broadens the ClpX mobility distribution

To test how heat shock affects ClpX mobility, we compared the D*_app_* distributions at 22 °C with those after 5 and 15 min at 32 °C, and after 5, 15 and 30 min of recovery at 22 °C (Fig. 3; Table 1). Heat shock broadened the distribution without significantly shifting its centre. After 5 min at 32 °C, the fractions of tracks below 0.10 µm²/s and above 0.60 µm²/s rose to 12.8% and 12.9%, while the median remained largely unchanged. After 15 min the broadening was more pronounced: a shoulder appeared at higher mobilities, the median decreased to 0.22 µm²/s and the low- and high-mobility fractions reached 14.8% and 14.2%. Consequently, the fraction of tracks in the intermediate range fell from 84.3% at 22 °C to 74.3% and 71.0% after 5 and 15 min at 32 °C and remained below the control value throughout recovery (77.8%, 75.5% and 78.4% after 5, 15 and 30 min).

**Figure 3.**
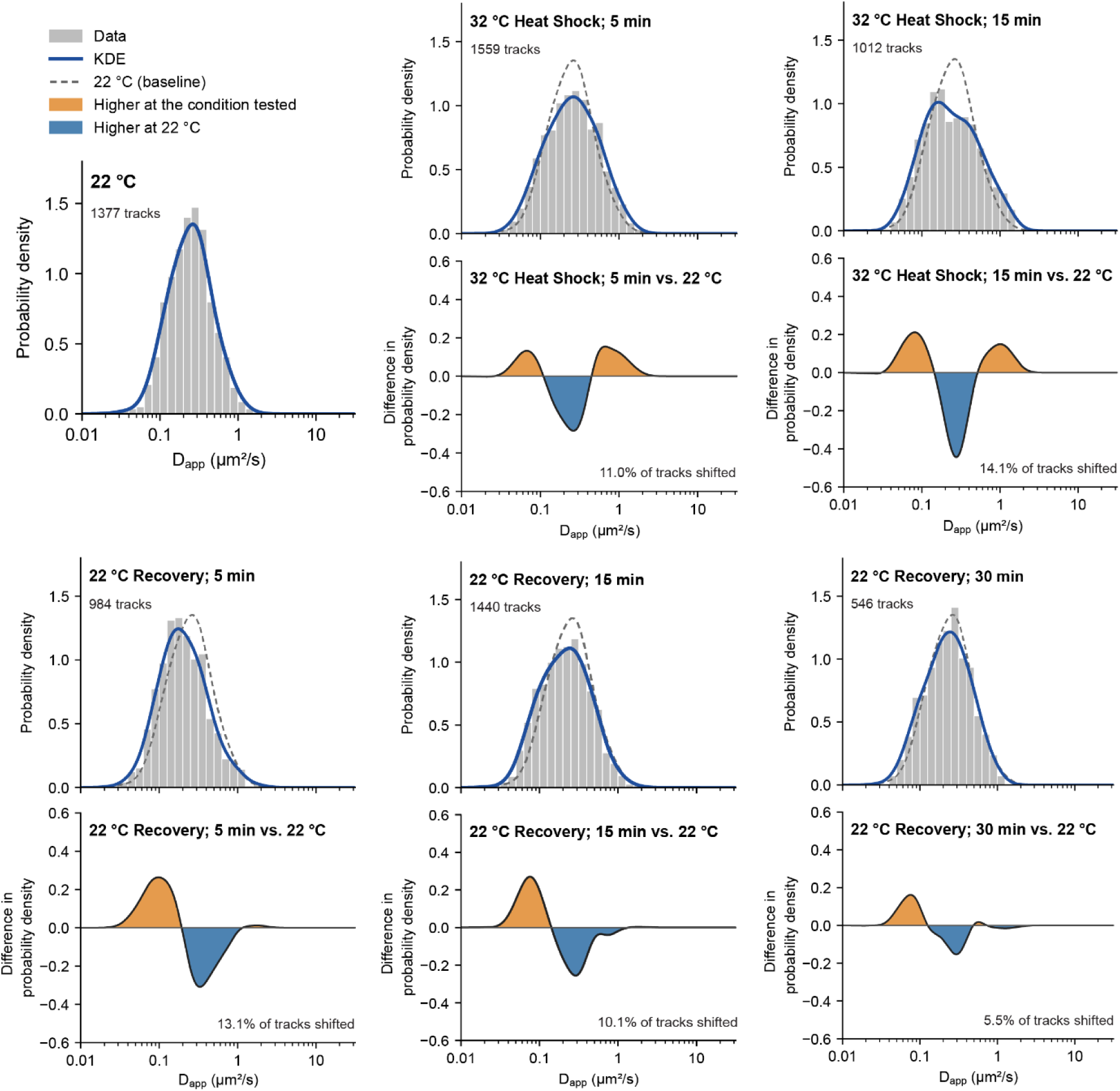
ClpX-HT mobility during heat shock and recovery. The figure shows D_app_ distributions (log-scale) of ClpX-HT tracks at 22 °C, after 5 and 15 min at 32 °C, and after 5, 15 and 30 min of recovery at 22 °C, and the respective difference between the kernel density estimate of each condition and that of 22 °C. Grey bars: probability density histograms; blue lines: Gaussian kernel density estimates (bandwidth 0.11 log₁₀ units); dashed grey lines: baseline at 22 °C. Orange: Dapp ranges more frequent in the condition; blue: more frequent at 22 °C. The percentage of tracks that shifted between D_app_ ranges relative to 22 °C is given in each panel. All data from three biological replicates.

**Table 1.**
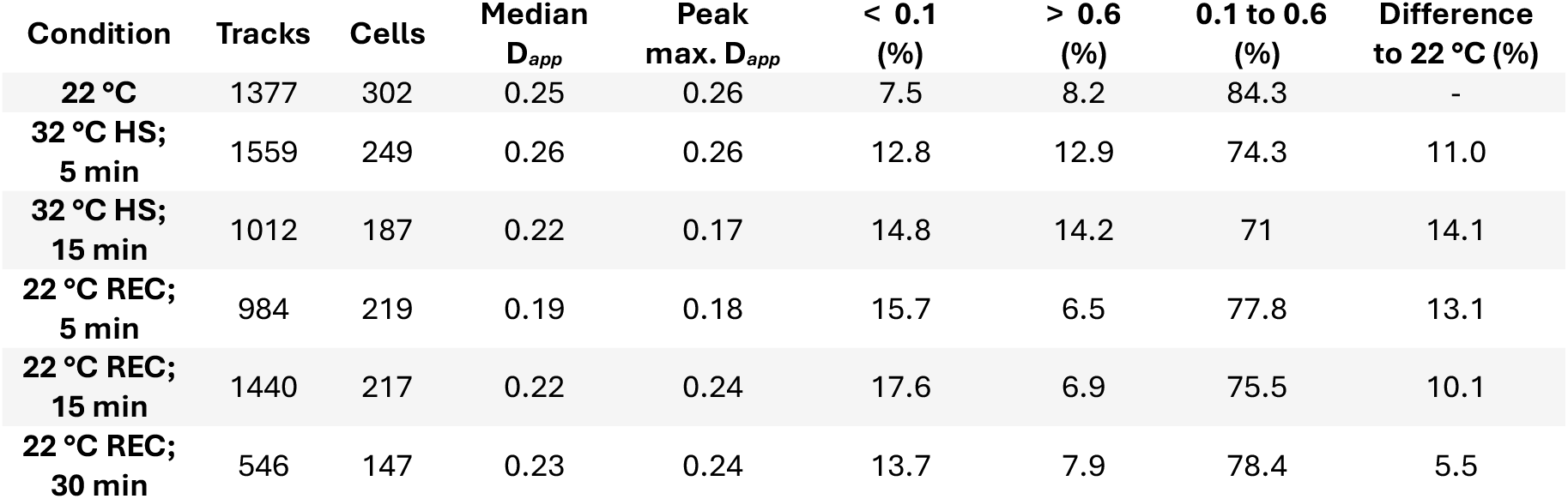
Summary statistics of ClpX-HT apparent diffusion coefficients. D*_app_* unit is µm²/s of individual trajectories. HS – heat shock, REC – recovery.

| Condition | Tracks | Cells | Median D<sub>*app*</sub> | Peak max. D<sub>*app*</sub> | < 0.1 (%) | > 0.6 (%) | 0.1 to 0.6 (%) | Difference to 22 °C (%) |
|---|---|---|---|---|---|---|---|---|
| **22 °C** | 1377 | 302 | 0.25 | 0.26 | 7.5 | 8.2 | 84.3 | - |
| **32 °C HS; 5 min** | 1559 | 249 | 0.26 | 0.26 | 12.8 | 12.9 | 74.3 | 11.0 |
| **32 °C HS; 15 min** | 1012 | 187 | 0.22 | 0.17 | 14.8 | 14.2 | 71 | 14.1 |
| **22 °C REC; 5 min** | 984 | 219 | 0.19 | 0.18 | 15.7 | 6.5 | 77.8 | 13.1 |
| **22 °C REC; 15 min** | 1440 | 217 | 0.22 | 0.24 | 17.6 | 6.9 | 75.5 | 10.1 |
| **22 °C REC; 30 min** | 546 | 147 | 0.23 | 0.24 | 13.7 | 7.9 | 78.4 | 5.5 |

The two tails recovered on different timescales. The fast fraction returned to control levels within 5 min at 22 °C (6.5% above 0.60 µm²/s, compared with 8.2% at 22 °C), while the slow fraction remained elevated (15.7% below 0.10 µm²/s after 5 min and 17.6% after 15 min). The distribution after 5 min of recovery was therefore shifted towards lower mobilities, with the lowest median of all conditions (0.19 µm²/s), while excess of slow tracks persisted at 15 min. After 30 min the distribution most closely resembled 22 °C: the median (0.23 µm²/s) and fast fraction (7.9%) matched 22 °C values, but the slow fraction remained nearly twice as high (13.7% versus 7.5%). Together, heat shock at 32 °C broadens the distribution of ClpX mobility, increasing both the slow and fast fractions. During recovery the fast fraction returns to baseline within 5 min, whereas the slow fraction persists for at least 30 min.

### Heat shock transiently redistributes ClpX to the cell periphery

At 22 °C, ClpX-HT localization was spread throughout the cytoplasm (Fig. 4a). Along the long axis, the distribution followed the homogeneous reference closely, with no clear enrichment at the poles. Across the short axis, the profile matched the expected profile for an even distribution in a rod-shaped cell. After 5 min at 32 °C, the long-axis distribution remained close to homogeneous, but the short-axis profile flattened, with more localizations towards the cell sides and fewer at the centre (Fig. 4b). After 15 min, the distribution was the least uniform of all conditions (Fig. 4c): localizations were enriched near the poles and the cell periphery, visible in the 2D map as discrete high-density spots.

**Figure 4.**
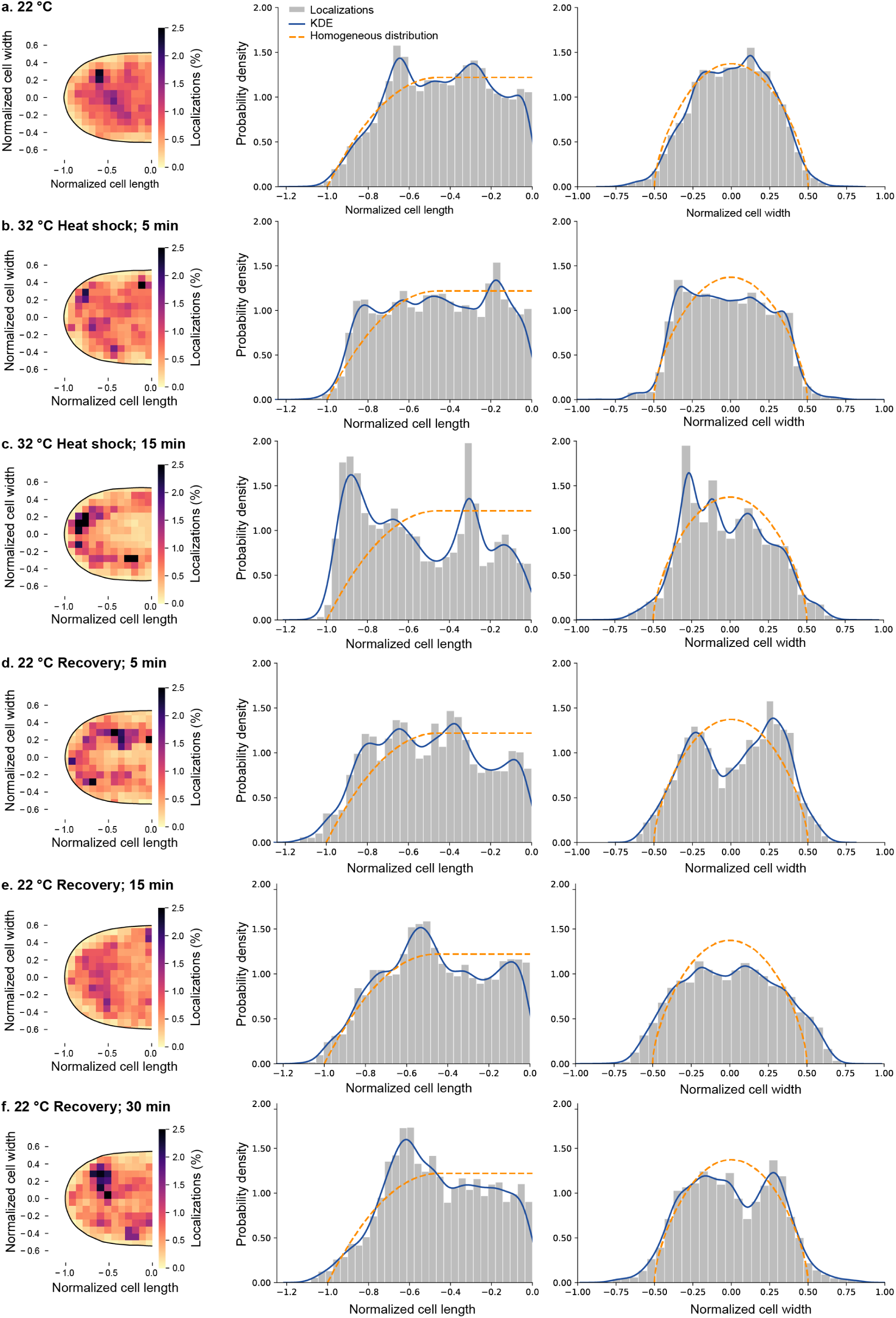
ClpX-HT localization during heat shock and recovery. Localizations of ClpX-HT molecules in P. cryohalolentis cells kept at 22 °C (**a**), heat-shocked at 32 °C for 5 min (**b**) and 15 min (**c**) and returned to 22 °C for 5 min (**d**), 15 min (**e**) and 30 min (**f**). Left: 2D localization maps showing all localizations for each condition in normalized, folded cell coordinates (pole at −1, mid-cell at 0). Colour palette shows the percentage of localizations per bin. Black line, cell outline. Middle and right: 1D distributions along the long axis (PC0, normalized length) and the short axis (PC1, normalized width). Gray bars, density histograms. Blue lines, KDE (Scott’s bandwidth × 0.C3). Orange dashed lines, the profile expected for molecules spread evenly through the volume of a rod-shaped cell (a cylinder with hemispherical caps), smoothed with the same kernel. All data from three biological replicates.

After 5 min of recovery at 22 °C, the long-axis distribution returned close to homogeneous while the short-axis profile became clearly bimodal (Fig. 4d), with a pronounced dip at the cell centre, indicating that ClpX was concentrated at the cell periphery rather than at the poles. This pattern disappeared with longer recovery: the central dip was shallower after 15 min (Fig. 4e), and by 30 min the short-axis profile again matched the homogeneous reference (Fig. 4f).

Together, heat shock and early recovery transiently redistribute ClpX. The change is mainly across the cell width, towards the periphery, and is most pronounced after 5 min of recovery. It largely reverts within 30 min at 22 °C, over the same time frame in which the D_app_ distribution returned towards the 22 °C profile (Fig. 3).

## Discussion

In this study, we established a protocol for single molecule tracking (SMT) in the psychrophilic bacterium *P. cryohalolentis*, based on an inducible expression system for HaloTag fusion proteins. To our knowledge, this is the first application of SMT in a psychrophile. Using the AAA+ unfoldase ClpX as a proof of concept, we show that the dynamics of individual proteins can be followed in a psychrophile *in vivo*, and that this approach detects changes in protein mobility in response to environmental stress. Studying this group of microorganisms at the molecular level *in vivo* has been technically challenging, and the platform presented here opens the possibility of studying how temperature affects their protein behaviour.

At 22 °C, ClpX-HT showed a unimodal distribution of apparent diffusion coefficients centred at 0.25 µm²/s. This remains below what would be expected for a freely diffusing particle the size of a ClpX hexamer (300–475 kDa with HaloTag), based on the size dependence of cytoplasmic diffusion in *E. coli* [41, 42]. It is instead comparable to values reported for free ribosomal subunits (∼1 MDa) [23, 43]. The low mobility therefore indicates that the fusion is incorporated into large complexes and suggests that a substantial fraction of ClpX is associated with ClpP (ClpXP, 0.7–1.2 MDa) and with substrates already under non-stress conditions, consistent with the role of ClpXP in protein turnover during normal growth [44]. Because the reference values were obtained in *E. coli*, this comparison is qualitative.

Heat shock at 32 °C broadened the D*_app_* distribution, increasing the proportions of both slow and fast molecules. The two populations behaved differently during recovery. While the fast fraction returned to control levels within 5 min, the slow fraction remained elevated for at least 30 min. This divergence suggests that they reflect distinct processes. ClpX participates in the heat-shock response in other bacteria [20, 19], and a greater demand for degradation of misfolded proteins during heat stress would be expected to increase the fraction of ClpX engaged with ClpP and substrates. The persistent slow population fits this interpretation, and its gradual return to baseline may reflect the time needed to clear proteins damaged during the shock.

During late heat shock and early recovery, ClpX localized to the cell periphery rather than to the poles. In *E. coli*, heat-induced protein aggregates accumulate predominantly at the cell poles [45], whereas large complexes such as translating ribosomes are excluded from the nucleoid and enriched at the cell periphery [23]. The peripheral pattern is therefore more consistent with large, mobile ClpX-containing complexes excluded from the nucleoid than with ClpX trapped in aggregates. Foci were occasionally observed, particularly 30 min into recovery, shifting the density towards the poles. ClpX and ClpP are known to form foci in several bacteria [35, 46, 47, 48], though these patterns may be organism-specific, reflecting differences in cell cycle regulation and in the available Clp partners.

The transient fast population may instead reflect partial, reversible disassembly of ClpX hexamers at elevated temperatures. Hexamer assembly depends on nucleotide binding [18, 49], and the subunit interfaces of a cold-adapted ClpX may be particularly sensitive above the growth optimum, since psychrophilic proteins are generally more flexible and less thermally stable than their mesophilic counterparts [14, 15]. Rapid reassembly on return to 22 °C would explain the disappearance of the fast fraction within the first minutes of recovery. A second, not mutually exclusive possibility is that ClpX is released from ClpP during heat shock because ClpP is recruited by competing Clp ATPases. In *B. subtilis*, heat shock reduced the number of ClpX foci while ClpP foci increased [35], probably reflecting ClpP association with ClpC and ClpE [44]. *P. cryohalolentis* lacks these proteins but encodes ClpA (NCBI RefSeq assembly: GCF_000013905.1), which has a different substrate specificity but, like ClpX, delivers substrates to degradation by ClpP [17, 50, 51]. Competition between ClpX and ClpA for ClpP under stress could free part of the ClpX pool and contribute to the fast population. A general change in cytoplasmic viscosity, which is temperature-dependent and sensitive to metabolic activity [52], is a less likely explanation, since it would shift the whole distribution rather than broaden it in both directions as observed here.

Clp proteases may be especially important for protein quality control in this organism. Many bacteria, including the psychrophile *Pseudoalteromonas translucida*, encode the Lon protease [53], a major contributor to the degradation of misfolded proteins [54, 55], but we did not identify a Lon homologue in the *P. cryohalolentis* genome (NCBI RefSeq assembly: GCF_000013905.1). If confirmed, ClpP-based proteases (ClpXP and ClpAP) would carry a larger share of the response to protein damage here, making *P. cryohalolentis* an interesting model for studying Clp proteases in cold adaptation and thermal stress.

Several limitations should be considered. Because only plasmid-encoded subunits carry HaloTag and the chromosomal *clpX* gene was retained, the tracked molecules represent a subset of the cellular pool assembled into mixed hexamers with native subunits. Relative fractions and distributions should be unaffected if labelling is random, but absolute numbers are underestimated. We did not test the activity of these mixed hexamers and complementation of a Δ*clpX* strain will be important for future quantitative work. For the same reason, the number of dye molecules per particle could not be determined, so fluorescence intensity alone cannot distinguish intact from partially disassembled hexamers, and the proposed disassembly remains to be tested directly. However, the low D*app* of ClpX-HT compared with free HaloTag (D*app* 5.6 µm²/s in *E. coli* [36]) indicates that the fusion is incorporated into large complexes, and its stress-dependent changes in mobility and localization show that it responds to heat shock. Moreover, tag-induced clustering is also a known artefact of fusion constructs: Clp proteases have previously been reported to form foci depending on the tag in *E. coli* [56], while HaloTag-ClpP forms highly mobile particles throughout the cytoplasm with only weak residual foci [57]. HaloTag fusions are less prone to this than many fluorescent proteins, but clustering cannot be fully excluded; the few cells containing immobile bright spots were omitted from our analysis.

Future work could track ClpP and ClpA under the same conditions, use free HaloTag as a direct mobility reference in *P. cryohalolentis*, and compare ClpX variants lacking ClpP docking [58] or ATP hydrolysis [18] activity. Beyond protein quality control, the expression system is a general SMT platform as almost any protein can be fused to HaloTag and tracked the same way, giving access to transcription, translation, DNA repair and cell division in a cold-adapted organism. The adaptations attributed to psychrophiles are claims about molecular behaviour in the cell, yet they have been inferred mostly from purified components or population averages [14, 59, 60]. Comparing the same process in a psychrophile and a mesophile across their physiological temperature ranges would test whether cold adaptation alters the kinetics of these processes or merely shifts the temperature at which they operate. Our results demonstrate that SMT can resolve stress-induced changes in protein dynamics in a psychrophilic bacterium, opening the way to studying molecular adaptation to cold *in vivo*.

## Data availability

Strains generated in this study are available upon request. The analysis codes are available at: https://github.com/Makelalab/Published/tree/main/Nery_et_al_2026

## Author contributions

Conceptualization, E.T.N. and J.M.; Methodology, E.T.N.; Investigation, E.T.N.; Funding acquisition, J.M.; Formal analysis, E.T.N. and Y.S.; Data curation, E.T.N.; Visualization, E.T.N.; Project administration, J.M; Software E.T.N. and Y.S.; Supervision, J.M.; Writing – original draft, E.T.N. and J.M.; and Writing – review C editing, E.T.N., Y.S., and J.M.

## Acknowledgements

We thank other members of the Mäkelä group for their support and discussions. The computations and data handling were enabled by resources provided by the CSC – IT Center for Science. This work was supported by the European Research Council (ERC StG TEMPADAPT, 101075984) and the Research Council of Finland (349116 and 353572). The funders had no role in study design, data collection and interpretation, or the decision to submit the work for publication.

## Supplementary Material

**Table S1.**
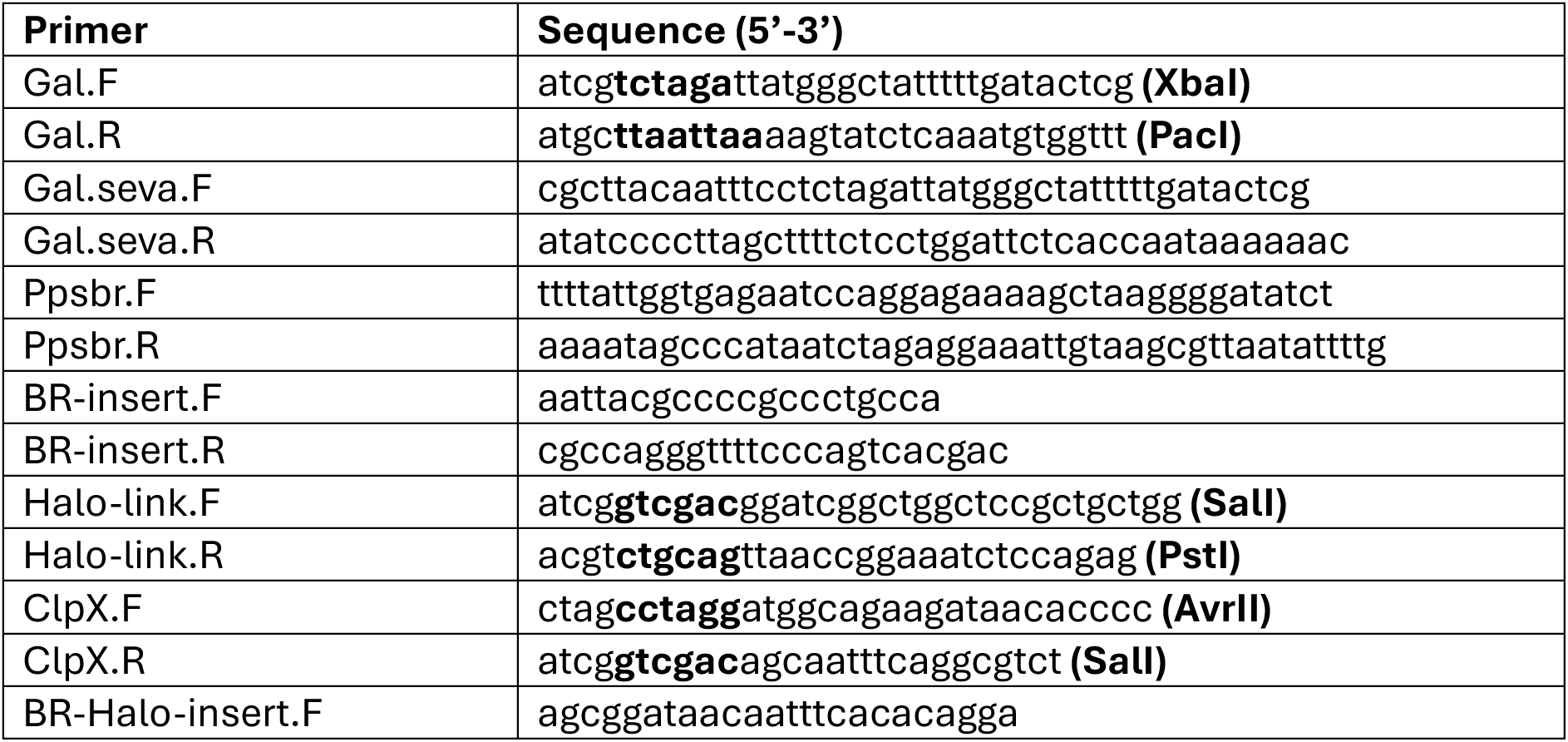
Table of primers.

**Figure S1.**
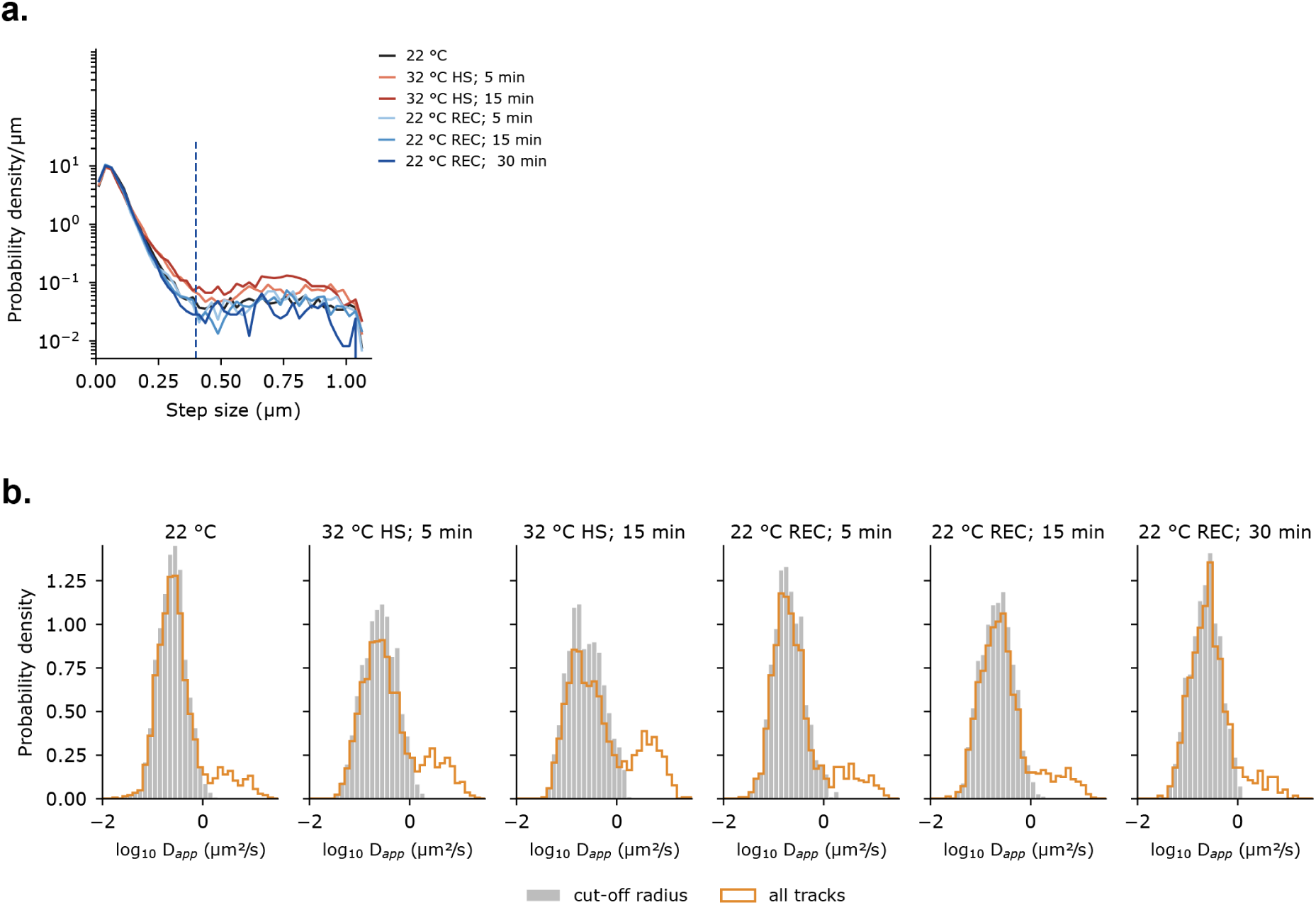
(**a.**) Distribution of step sizes between consecutive localizations for all conditions (HS – heat shock; REC – recovery). The dashed line marks the 0.4 µm cutoff; the dotted line marks the larger linking radius of 1.0C µm. (**b.**) Distributions of apparent diffusion coefficients for original tracks (orange) and after splitting (grey).

